# A unified neurocognitive model of the anterior temporal lobe contributions to semantics, language, social behaviour & face recognition

**DOI:** 10.1101/725515

**Authors:** Junhua Ding, Keliang Chen, Haoming Liu, Lin Huang, Yan Chen, Yingru Lv, Qing Yang, Qihao Guo, Zaizhu Han, Matthew. A. Lambon Ralph

**Author notes:** Joint corresponding authors: Qihao Guo: email -, Zaizhu Han: email -, Matt Lambon Ralph: email -, MRC Cognition and Brain Sciences Unit, University of Cambridge, 15 Chaucer Road, Cambridge, CB2 7EF UK. joint first author (Junhua Ding, Keliang Chen).

## Abstract

The anterior temporal lobes (ATL) have become a key brain region of interest in cognitive and clinical neuroscience. Contemporary explorations are founded upon neuropsychological investigations of semantic dementia (SD) that describe the patients’ selective semantic impairment and the variations in their language, behavioural and face recognition abilities. The purpose of this investigation was to generate a single unified model which captures the known cognitive-behavioural variations in SD, and integrates with the considerable database on healthy semantic function and other patient groups. A new analytical approach was able to capture the graded neuropsychological differences and map these to the patients’ distribution of frontotemporal atrophy. Multiple regression and principal component analyses confirmed that the degree of generalised semantic impairment was related to the patients’ total, bilateral ATL atrophy. Verbal production and word-finding abilities were related to total ATL atrophy as well as to the balance of left>right ATL atrophy. Behavioural apathy was found to relate positively to the degree of orbitofrontal atrophy and negatively to total temporal volumes. Disinhibited behaviour was related to right ATL and orbitofrontal atrophy and face recognition to right ATL volumes. Rather than positing mutually-exclusive sub-categories, the data-driven model repositions semantics, language, social behaviour and face recognition into a continuous frontotemporal neurocognitive space.

## Introduction

Recent years have shown a considerable increase of interest in the cognitive and behavioural functions of the anterior temporal lobes (ATL). This heightened attention on the ATL is founded upon the unique insights that arise from detailed neuropsychological and neurological investigations of semantic dementia (SD; the temporal-variant of frontotemporal dementia)^1–5^. These findings have, in turn, inspired cognitive neuroscience explorations of the contribution of the ATL to semantic representation in healthy participants^6–10^, comparative studies across different patient groups with impaired semantic performance^11–16^ and formal neuroanatomically-constrained computational models of semantic cognition^17–21^. The purpose of this investigation was to go beyond these descriptions to generate, for the first time, a unified neurocognitive model of ATL function which captures the known cognitive-behavioural variations across SD, maps these to the underlying patterns of atrophy, and integrates with the considerable database on the graded bilateral ATL contributions to healthy semantic function^22^ and semantic impairment in other patient groups^23^. In doing so, this approach accommodates the facts that: (a) there are not mutually-exclusive subtypes of SD but rather *gradely-varying* patterns of cognitive-behavioural presentation; (b) SD is a part of the frontotemporal dementia (FTD) spectrum; (c) the cognitive-behavioural deficits reflect not only the balance of left vs. right ATL atrophy but also the total temporal lobe atrophy and its extension to insular and orbitofrontal cortex (OFC); and (d) semantic representation in healthy participants appears to be supported by ATL regions, bilaterally, with graded variations in function reflecting differential patterns of connectivity.

The existing literature on SD and the associated explorations of ATL function in healthy participants and other patient groups, can be clustered around three descriptive ‘subtypes’. The principal description is of the cognitively-selective yet generalised degradation of concepts found in SD^1–3^. Thus, whilst many non-semantic aspects of higher cognition and perception can be strikingly preserved until relatively late, SD patients exhibit an insidious degradation of semantic representations. There is clear evidence that SD affects all categories of concept (living, non-living, concrete, abstract, nouns and verbs)^24–26^ in both receptive and expressive tasks for verbal and nonverbal domains^27–32^, with performance strongly influenced by concept familiarity, typicality and specificity^1, 33, 34^. It was this pervasive multimodal degradation which led Snowden and Neary to coin the term “semantic dementia” in 1989^2^. The more contemporary consensus criteria^35^ proposed the alternative term “semantic-variant primary progressive aphasia (PPA)” thereby contrasting it from progressive non-fluent aphasia and logopenic progressive aphasia. Whilst the patients’ pronounced anomia and verbal comprehension deficit are often prominent clinical features in the initial clinical presentation^5, 36^, careful evaluation invariably identifies a range of nonverbal comprehension deficits^29–32, 37^ even in early cases^28^. This striking clinical presentation is underpinned by bilateral atrophy centred on the polar and ventral aspects of the ATL^38, 39^. Patients with more left than right (L>R) ATL atrophy are typically the more common in most clinics^40^. Mild patients can present with predominantly left ATL atrophy or, more rarely, with right ATL atrophy (see below); however, the disease is inherently bilateral in nature such that cases with asymmetric atrophy are associated with considerable FDG-PET hypometabolism and rapid subsequent atrophy in the contralateral ATL^41, 42^. The combination of multimodal semantic impairment with bilateral ATL atrophy is consistent with the hypothesis that semantic memory is underpinned by a bilaterally-distributed representational system^20, 23^. This notion is also supported by data from: (i) patients with unilateral ATL damage (who have mild semantic impairment after left or right resection)^15^; (ii) bilateral ATL resection in non-human primates^43, 44^ and a single-case human study^45^ – in which initial unilateral resection generated a mild multimodal semantic impairment and then subsequent contralateral ATL resection led to a profound deficit; (iii) results from ATL repetitive transcranial magnetic stimulation (rTMS) in healthy participants (which generates a transient, selective multimodal semantic deficit after left or right ATL stimulation)^46^ and, (iv) fMRI investigations of semantic processing in healthy participants using methods that correct or minimise ATL-related distortion artefacts and other methodological limitations (which show bilateral ATL engagement by semantic tasks particularly in ventral and lateral regions)^8, 22, 47^.

A second related SD sample is of those patients who present early with predominantly left ATL atrophy. The clinical presentation is dominated by the patients’ profound anomia accompanied by a mild comprehension deficit^17, 36^. Again this pattern in SD has direct parallels in other patient groups and in healthy participants: (i) patients with left ATL damage/resection are significantly more anomic than their right counterparts^15, 48–50^; (ii) fMRI studies involving speech production (covert or overt) lead to greater left than right ATL activations^22^; and (iii) rTMS has a larger effect on picture naming after left than right ATL stimulation^51^. Two hypotheses have been proposed to explain these data: (a) the left ATL houses lexical representations that support semantically-driven speech production^48, 49^; or (b) that the bilateral ATL-hub semantic system connects to left-lateralised prefrontal speech production systems from the left ATL^17, 20^. Whilst both theories explain the differential anomia of left-ATL patients, the second approach is able to accommodate a range of additional findings, including some of those already noted above: (i) there are graded rather than absolute differences between left and right ATL cases/function; (ii) right unilateral ATL patients have some degree of anomia; (iii) careful examination of early L>R SD cases reveals mild deficits in verbal and nonverbal comprehension; (iv) the left ATL is activated in healthy participants for the same types of verbal and nonverbal comprehension tasks, and are compromised after rTMS.

The third SD sample relates to those patients with greater right ATL atrophy. Early presentation with predominantly right ATL atrophy is the least common in most clinics though the literature contains some notable single case studies and small case-series^52–58^. Studies have tended to focus on one of two phenomena. Many of the very early, predominantly right ATL cases show visual recognition deficits for familiar people, followed with progression (see the final section of the Results) to anomia, generalised multimodal person semantic deficits and ultimately the generalised semantic impairment associated with SD^53, 55^. A second sub-literature relates to the social and behavioural impairments observed in SD, which have been associated more with the R>L patients^40, 58–61^. An important question to resolve is how face recognition deficits and social-behavioural impairments fit with the other aspects of SD. Indeed, reaching a correct interpretation and unified model of SD is challenged by three facts: (a) the L>R patients also have behavioural changes when this is formally assessed^60^ as well as poor semantic knowledge about people (which might reflect the longstanding and well-established fact that specific-level concepts are more vulnerable to semantic degradation as observed in patients and formal computational models)^1, 18, 34^; (b) healthy participants activate superior and ventral ATL regions, bilaterally, when making social concept judgements^62–64^ and exhibit transient impairments when the left or right ATL is stimulated^65–67^; and (c) R>L patients atrophy tend to have more atrophy overall not only in bilateral temporal regions but also extending to OFC^60, 61, 68, 69^, which is known to be a behavioural-related region^70, 71^. Even one seminal study that matched pairs of left- and right-dominant cases for overall temporal atrophy found remaining differences in frontal atrophy^72^.

The aim of the current study, therefore, was to generate a data-driven, joint cognitive-neuroanatomical framework which assimilates these three clinical variations into a unified model and to test if the results align with those from contrastive patients (particularly those with unilateral ATL damage) and healthy participants. A recent computationally-informed theory suggests that semantic representations are supported by a bilateral ATL system which engages with multiple, distributed sources of verbal and sensory engrams to form coherent, generalizable concepts^23, 50^. Graded variations in function both within (e.g., dorsal vs. ventral) and between (left vs. right) the ATLs emerge as a natural consequence of differential connectivity to input and output systems^17, 20, 22, 73, 74^. For example, the heavy engagement of the left ATL in speech production^15, 49, 51, 75^ follows from connectivity to the strongly left-lateralised speech production systems^76^.

The unified model was generated in two steps. The first was to use multiple regression to explore the relationship between the primary symptoms of SD and the integrity of key frontotemporal regions. The use of multiple regression allows for shared variance in the distribution of atrophy and behaviours across patients to be modelled and separated. In the second step we utilised a data-driven approach – namely, to use principal component analysis (PCA) with varimax rotation to explore the underlying, graded dimensions of variation in the patients’ neuropsychological and social-behavioural data. This approach dispenses with a search for mutually-exclusive categories (which may not exist) and, instead, assumes that the patients vary in graded ways across one or more underlying dimensions^70, 77, 78^. Then the factor scores (rather than individual test scores) are related to the patients’ pattern of underpinning atrophy (whole-brain analyses). This PCA plus voxel-symptom mapping has been successfully applied by multiple independent research groups to post-stroke aphasia data, leading to consistent patterns of behavioural dimensions and neural correlates across different patient samples and assessment batteries^77–82^ as well as one recent exploration of impulsivity and apathy in frontotemporal lobar syndromes^70^. Accordingly, we extended the same methodology to a new large SD sample containing variation of left-right ATL atrophy and disease severity.

## Results

### The demographic, neuropsychological and behavioural profiles of SD patients

Table 1 summarises the demographic, neuropsychological and behavioural information pertaining to the SD patients. Patients were matched to the healthy controls in terms of age, gender and education years (*p* values > 0.07). As expected, SD patients presented with impairments of object and face semantics, visual perception and general cognitive ability (*p* values < 0.001) except for the preservation of visual object perception of right SD patients (*p* = 0.07). Additionally, some behavioural problems were also reported. The most six frequent symptoms included apathy (70%), irritability (59%), agitation (56%), anxiety (56%), depression (52%), and disinhibition (44%). With regard to cognitive and behavioural differences between left and right SD patients, left SD patients performed better on word picture verification of faces (*p* = 0.009), while right SD patients had better performance on visual object perception and picture naming of objects (*p* values < 0.002).

**Table 1.** Demographic characteristics, cognitive and behavioural performance and cerebral grey matter volumes of SD patients and healthy subjects. The numbers in parentheses are the standard deviations.

|  |  | Healthy controls<br>(n = 20) | SD patients |  |
| --- | --- | --- | --- | --- |
|  |  |  | left SD (n = 28) | right SD (n = 19) |
| **<u>Demographic characteristics</u>** |  |  |  |  |
| Age (years) |  | 61 (4) | 62 (7) | 63 (7) |
| Gender (male: female) |  | 8:12 | 16:12 | 9:10 |
| Education level (years) |  | 10 (3) | 12 (3) | 11 (3) |
| **<u>Behavioural performance</u>** |  |  |  |  |
| *Object semantics* | Oral picture naming | 92% (4%) | 30% (19%)* | 46% (20%)¶§ |
|  | Word picture verification | 96% (3%) | 67% (21%)* | 71% (20%)¶ |
|  | Picture associative matching | 95% (4%) | 78% (11%)* | 77% (8%)¶ |
| *Face semantics* | Oral picture naming | 78% (16%) | 10% (9%)* | 13% (13%)¶ |
|  | Word picture verification | 93% (9%) | 54% (22%)* | 38% (22%)¶§ |
|  | Picture associative matching | 96% (6%) | 72% (17%)* | 69% (14%)¶ |
| *Visual perception* | Object perception | 96% (5%) | 72% (20%)* | 87% (13%)§ |
|  | Face perception | 78% (14%) | 58% (18%)* | 54% (18%)¶ |
| *General cognition* | MMSE | 28 (1) | 22 (4)* | 23 (4)¶ |
| *NPI (0:1:2:3)* | Agitation |  | 7:4:1:1 | 5:5:3:1 |
|  | Depression |  | 8:2:2:1 | 5:5:1:3 |
|  | Anxiety |  | 6:3:4:0 | 6:7:0:1 |
|  | Apathy |  | 5:5:3:0 | 3:7:3:1 |
|  | Disinhibition |  | 9:2:1:1 | 6:6:1:1 |
|  | Irritability |  | 6:4:3:0 | 5:4:3:2 |
| **<u>Cerebral volume</u>** |  |  |  |  |
| Total grey matter volume (mm<sup>3</sup>) |  | 32156 (2496) | 29522 (3508)* | 28319 (3624)¶ |
| Left – right ATL volume (mm<sup>3</sup>) |  | -10 (43) | -191 (65)* | 167 (82)¶§ |
| Left + right ATL volume (mm<sup>3</sup>) |  | 1625 (163) | 1092 (238)* | 942 (148)¶§ |
| Left + right OFC volume (mm<sup>3</sup>) |  | 1144 (94) | 1002 (122)* | 897 (155)¶§ |

\* indicates P values of left SD vs controls < 0.05; ¶ indicates P values of right SD vs controls < 0.05; § indicates P values of left vs right SD < 0.05. SD: semantic dementia; MMSE: Mini Mental State Examination; NPI: neuropsychiatric inventory.

### The atrophy of SD patients

The distribution and balance of atrophy in this SD sample was typical of those reported by other research groups. Specifically, the voxel-based analysis revealed that both left and right SD patients had marked atrophy in the bilateral temporal lobes, insula and OFC (see Figure 1a & b). When directly comparing the two SD subgroups, right SD showed more atrophy in the right temporal lobe, insula and OFC, while additional atrophy in the left SD subgroup was mainly restricted to the left temporal lobe (see Figure 1c).

**Figure 1.**
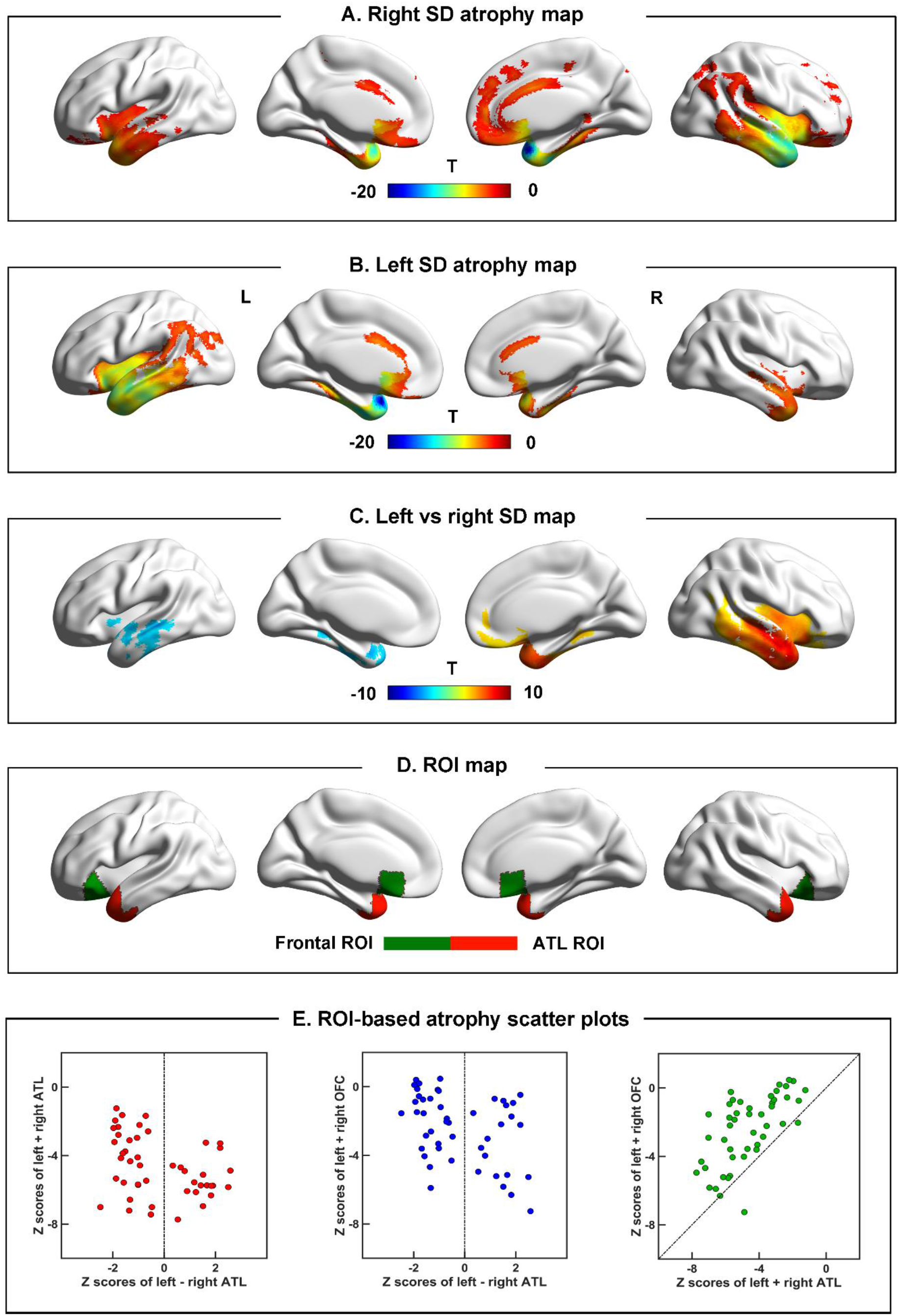
The atrophy pattern of SD patients. Panel A & B: the comparison between left SD vs. controls and right SD vs. controls (FDR corrected *p* < 0.05). Panel C: the comparison between left and right SD (FDR corrected *p* < 0.05). Panel D: the ROIs used in the behavioural-atrophy regression analyses. Panel E: scatter plots showing the distribution and inter-relationships of left ATL, right ATL and OFC atrophy across the SD patient sample. All the brain maps were made using Brainnet^83^. ATL: anterior temporal lobe; OFC: orbital frontal cortex; SD: semantic dementia.

As noted in the Introduction, simple comparison of left and right SD patients is complicated by the fact that other potentially important differences occur across the groups. In keeping with previous reports^60, 68^, the ROI analysis confirmed: (1) atrophy for the SD patient group as a whole was found in bilateral ATL and OFC relative to controls (*p* values < 0.001; see Table 1); and (2) comparison between left and right SD revealed that the right SD patients had more atrophy to both ATLs (*p* = 0.01) and OFC (*p* = 0.006; see Table 1). This is important for interpretation of any cognitive or behavioural differences between left and right SD patients for two reasons: (A) the right SD patients were neuroanatomically more severe and (B) their atrophy also encroached into potentially important regions such as the left ATL and OFC. These relationships can be clearly observed in the scatterplots shown in Figure 1e: (i) there is no boundary between left and right SD cases; (ii) most patients have a degree of bilateral ATL and OFC damage, with the exception of some very mild left SD cases (indeed, OFC damage is correlated with total temporal damage (see Figure 1e, right panel) though, as expected for an SD sample, there is always greater temporal than OFC atrophy; replicating the results from Seeley et al^72^); (iii) the right SD cases have greater bilateral ATL (Figure 1e, left panel) and OFC (Figure 1e, middle panel) damage; and (iv) very mild right-ATL only cases are extremely rare (indeed, like many group SD studies, no such cases were found in our clinical sample; thus there are no cases in the upper right-hand region of the left scatterplot in Figure 1e). Accordingly, rather than using a simple left-right binary division of SD cases, we adopted new statistical approaches which take into account not only the laterality but also total temporal and OFC atrophy. In the first step we used simultaneous regression (to map these anatomical measures to the cognitive and behavioural results) and then we utilised PCA to establish the underlying dimensions of behavioural variation in the SD sample and explored their relationship to the atrophy distributions.

### The relationships between atrophy measures and task performance

Simultaneous regression analyses were conducted using the three ROI atrophy measures to predict each task. As shown in Table 2, for two word-picture verification tasks, the sum of ATL volume was the only significant variable (*b* values > 0.42, *p* values < 0.02). Two picture naming tasks were also predicted by the sum of ATL volume (*b* values > 0.44, *p* values < 0.02) as well as the difference of left>right atrophy (*b* values > 0.35, *p* values < 0.02). For two perception tasks, the object perception was related with the ATL atrophy difference (*b* = 0.35, *p* = 0.05), while the face perception performance was related with total OFC atrophy (*b* = 0.48, *p* = 0.02). Regarding the NPI measures, only anxiety and apathy showed significant relationship with the atrophy measures (it may be important to note that agitation, depression, disinhibition and irritability were rated as 0 or 1 for the vast majority of the SD participants and thus there is a ceiling effect for these behavioural measures). Anxiety levels exhibited a significant *negative* relationship (*b* = −0.49, *p* = 0.03) with total ATL atrophy (perhaps reflecting heightened anxiety with the onset and diagnosis of the disease). Apathy was positively related to OFC atrophy (*b* = 0.59, *p* = 0.01) yet negatively with the sum of ATL (*b* = −0.48, *p* = 0.03). This would imply that, in the context of SD being a more temporal than frontal variant frontotemporal dementia, those cases with more balanced OFC and ATL atrophy are the most likely to demonstrate apathy.

**Table 2.** B values of 3 ROI variables predicted behavioural variables.

|  |  | Left – right<br>ATL volume | Left + right<br>ATL volume | Left + right<br>OFC volume |
| --- | --- | --- | --- | --- |
| **Face semantics** | Oral picture naming | 0.35* | 0.44* | -0.02 |
|  | Word picture verification | -0.09 | 0.42* | 0.04 |
|  | Picture associative matching | 0.09 | 0.35 | 0.15 |
| **Object semantics** | Oral picture naming | 0.55*** | 0.49** | -0.18 |
|  | Word picture verification | 0.29 | 0.53** | -0.14 |
|  | Picture associative matching | -0.07 | 0.21 | 0.13 |
| **Visual perception** | Object perception | 0.35* | -0.06 | 0.18 |
|  | Face perception | -0.09 | -0.09 | 0.48* |
| **NPI** | Agitation | -0.13 | -0.15 | 0.27 |
|  | Depression | -0.14 | -0.27 | -0.08 |
|  | Anxiety | 0.06 | -0.49* | 0.00 |
|  | Apathy | -0.07 | -0.48* | 0.59** |
|  | Disinhibition | -0.28 | -0.25 | -0.01 |
|  | irritability | -0.11 | -0.13 | -0.05 |
| **PCA** | Factor 1: apathy | 0.20 | -0.08 | 0.18 |
|  | Factor 2: face | -0.17 | 0.21 | 0.33 |
|  | Factor 3: naming | 0.62*** | 0.53*** | -0.28 |
|  | Factor 4: disinhibition | -0.14 | -0.04 | 0.09 |
|  | Factor 5: SD severity | 0.18 | -0.18 | 0.48* |

\* $p < 0.05$ ; \*\* $p < 0.01$ ; \*\*\* $p < 0.001$ .
ATL: anterior temporal lobe; OFC: orbital frontal cortex; NPI: neuropsychiatric inventory;
PCA: principal component analysis.

### PCA of task performance

All neuropsychological and behavioural data were entered into a PCA. The KMO for the resultant model was 0.71. Five factors were found as the optimal number of our data, then the missing data were imputed by probabilistic PCA. Using the imputed data, PCA were performed with varimax rotation. Figure 2 (top panels) displays the loadings of the tasks on each of the orthogonal factors. Five factors with an eigenvalue>1 were extracted, which accounted for 85% of the total variance. Factor 1 accounted for 21% of the variance. The loadings on Factor 1 were high for apathy (0.74), depression (0.89) and anxiety (0.72), and thus we refer to this factor as “apathy”. Factor 2 accounted for 20% of the variance. High loadings on Factor 2 were found for the visual aspects of face recognition (rather than semantics more generally) in the form of face matching (0.84) and face verification (0.84) – thus we refer to this as the “face” factor. Factor 3 (variance = 16%) was interpreted as ‘naming’ because it heavily loaded on three verbal tasks (face naming: 0.75; object naming: 0.90; object verification: 0.69). Factor 4 (variance = 16%) was labelled ‘disinhibition’ due to its high loadings with disinhibition (0.89), agitation (0.88) and irritability (0.65). Factor 5 accounted for 12% of variance, which had high positive loadings on object (0.86) and face (0.77) perceptions. We refer to this factor as ‘SD severity’.

**Figure 2.**
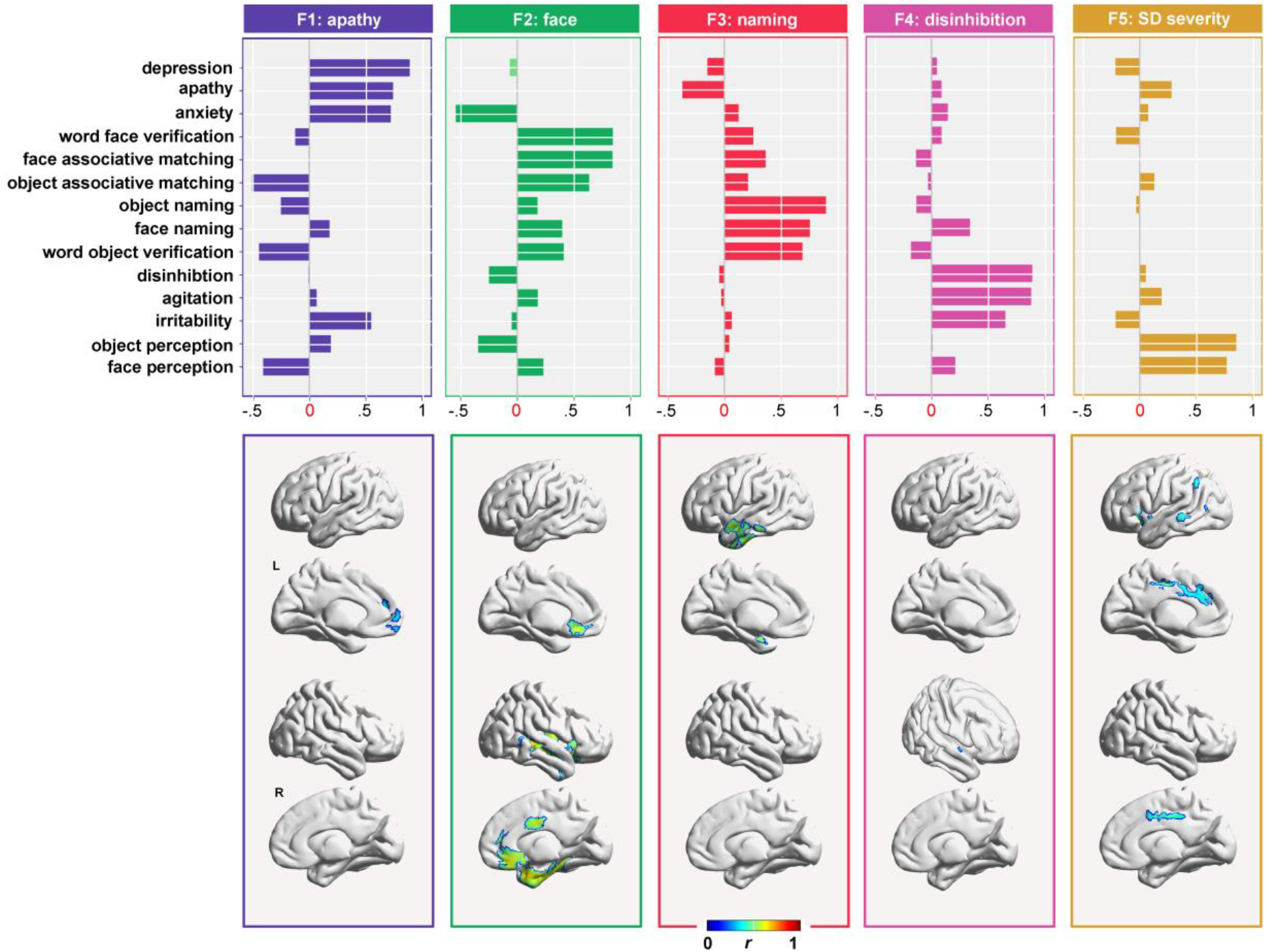
The factor loadings of PCA and corresponding maps of the significant regions associated with each factor (voxel size > 50).

### The relationship between atrophy and PCA factors

In the first symptom-atrophy mapping analysis, we used the three ROI variables in a multiple regression to predict the five PCA scores (as had been done for the individual test scores, see above). As shown in Table 2, the factor ‘SD severity’ were predicted by the sum of OFC volume (*b* value = 0.48, *p* values = 0.02). The sum and difference of bilateral ATL volumes showed significant effects for the ‘naming’ factor (*b* values > 0.53, *p* values < 0.001). None of the ROI variables was significantly related to the ‘disinhibition’, ‘apathy’ and ‘face’ factors.

Rather than limit the symptom-atrophy mapping to the ROI regions alone, in the second investigation we utilised voxel-based correlation mapping (VBCM) to provide a whole-brain analysis. These results replicated those found in the ROI-based analysis but also revealed additional areas of interest (see the lower panels in Figure 2 & Table 3). The SD severity factor was correlated with bilateral middle cingulate gyrus, left posterior temporal and parietal regions. Unsurprisingly, these regions represent the edges of the atrophy distribution in SD (see Figure 1) and the progression of atrophy as observed in previous longitudinal studies^42, 84, 85^. The naming factor was positively related to the atrophy of the left ATL. The right dorsal superior temporal gyrus (STG) and rectus gyrus were the only regions to be related to the inhibition factor. The apathy factor was associated with the atrophy of bilateral medial frontal cortex only. Finally, the face recognition factor was related to various areas within the right temporal lobe, insula and bilateral medial frontal lobes.

**Table 3.** The peak information of voxel-wised PCA correlation analysis

|  |  | Coordinates | R values | Voxel number | Region |
| --- | --- | --- | --- | --- | --- |
| **Factor 1: apathy**<br>($p < 0.05$) | Cluster 1 | 3 61.5 -12 | 0.38 | 111 | Right medial orbital frontal and rectus gyri |
|  | Cluster 2 | -4.5 61.5 -12 | 0.37 | 73 | Left medial orbital frontal and rectus gyri |
|  | Cluster 3 | -4.5 61.5 4.5 | 0.40 | 141 | Left superior medial and medial orbital gyri |
|  | Cluster 4 | -6 48 22.5 | 0.36 | 140 | Left superior medial and anterior cingulate gyri |
| **Factor 2: face**<br>($p < 0.001$) | Cluster 1 | 55.5 -28.5 -6 | 0.76 | 13315 | Right insula, middle temporal and temporal pole gyri |
|  | Cluster 2 | -1.5 31.5 -16.5 | 0.60 | 438 | Left rectus and medial orbital frontal gyri |
|  | Cluster 3 | 4.5 -3 34.5 | 0.60 | 397 | Right middle cingulate gyrus |
| **Factor 3: naming**<br>($p < 0.001$) | Cluster 1 | -37.5 -16.5 -37.5 | 0.71 | 4348 | Left inferior and middle temporal gyri |
|  | Cluster 2 | -21 3 -25.5 | 0.59 | 762 | Left temporal pole and parahippocampal gyri |
| **Factor 4: disinhibition**<br>($p < 0.05$) | Cluster 1 | 3 13.5 -24 | 0.35 | 76 | Right rectus and olfactory gyri |
|  | Cluster 2 | 51 6 -12 | 0.32 | 83 | Right temporal pole |
|  | Cluster 3 | 58.5 3 -1.5 | 0.35 | 52 | Right temporal pole |
| **Factor 5: SD severity**<br>($p < 0.01$) | Cluster 1 | -67.5 -33 -4.5 | 0.46 | 300 | Left middle temporal gyrus |
|  | Cluster 2 | -33 18 -10.5 | 0.52 | 484 | Left insula and inferior frontal gyri |
|  | Cluster 3 | -40.5 -15 -1.5 | 0.43 | 109 | Left insula and superior temporal gyri |
|  | Cluster 4 | -58.5 -69 -3 | 0.42 | 112 | Left middle temporal gyrus |
|  | Cluster 5 | -37.5 1.5 10.5 | 0.43 | 103 | Left insula gyrus |
|  | Cluster 6 | -1.5 -15 42 | 0.52 | 866 | Left anterior and middle cingulate gyri |
|  | Cluster 7 | 1.5 40.5 22.5 | 0.42 | 78 | Left and right anterior cingulate gyri |
|  | Cluster 8 | 1.5 -4.5 48 | 0.53 | 615 | Right middle cingulate and supplementary motor gyri |
|  | Cluster 9 | -49.5 -48 45 | 0.51 | 191 | Left inferior parietal gyrus |

2 Only positive clusters were reported. Cluster size > 50 voxels.

### Supplementary analysis of face-related ROIs in SD face processing deficits

Given that the PCA-VBCM highlighted various right temporal areas beyond the ATL, we explored the relationship of atrophy in five face-related ROIs (derived from studies of healthy participants) and the face-related PCA factor. Atrophy in all five face-related ROIs were highly correlated with the face factor (*r* values > 0.37, *p* values < 0.02; see Figure 3). Further validation analyses are reported in the supplementary materials, which revealed that this result for face ROIs was specific to face but not object processing, specific to SD patients but not normal controls, and only to right-sided but not the corresponding left homologues.

**Figure 3.**
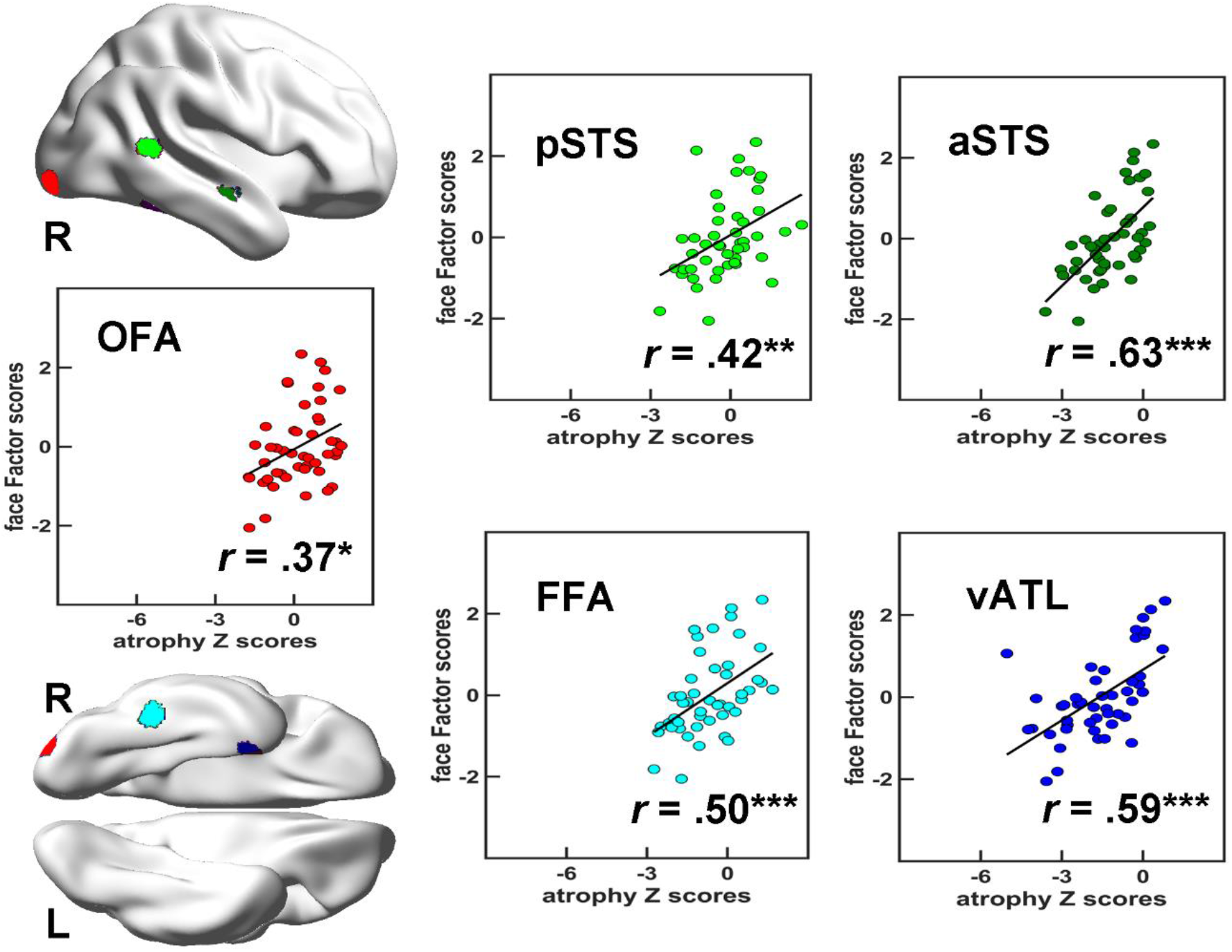
Correlation results between face-related ROIs and the PCA scores. The brain maps show the locations of ROIs. *: *p* < 0.05; **: *p* < 0.01; ***: *p* < 0.01. OFA: occipital face area; FFA: fusiform face area; pSTS: posterior superior temporal sulcus; aSTS: anterior superior temporal sulcus; vATL: ventral anterior temporal lobe.

### Testing the generalisation of the PCA-VBCM model: extrapolation to early predominantly right-sided SD

The preceding regression analyses and PCA-VBCM indicate that SD patients can be conceptualised within a unified multidimensional model. If correct then it should be possible not only to interpolate within the existing data but, more challengingly, to extrapolate to regions of the space for which there were no data. As noted above, like many other SD group samples, there were no patients with very mild predominantly right ATL atrophy. Extrapolating from the existing model, the presentation of very early right ATL-only patients should be dominated by the right-ATL loading Factor 2 & 4, namely visual face recognition and positive behavioural deficits. Then, longitudinally, not only would there be augmented atrophy in the right ATL but it should also become more bilateral in nature. Ultimately, atrophy should also encroach on the insula and OFC. Accordingly, the initial face recognition and positive behavioural deficits should be joined next by generalised semantic and naming impairments (associated with a more bilateral ATL picture) and then with negative behavioural features (associated with the OFC atrophy).

To test this hypothesis we conducted a systematic review of the literature, selecting case reports of patients who presented with early right predominant ATL atrophy. In particular, we focussed on investigations that provided detailed neuropsychological evaluation of the factors of interest (face recognition, naming, semantic abilities and behavioural impairment) as well as information about how these progressed over time (with respect to the patient’s history prior to clinical presentation or through systematic repeated assessment over time). We identified 15 cases in the literature and a summary of their symptom progression is provided in Table 4. With one exceptions, all other cases reflected the predicted neuropsychological pattern; namely, patients presented first with early visual face recognition or positive behavioural deficits followed by more general semantic, naming and negative behavioural problems.

**Table 4.**
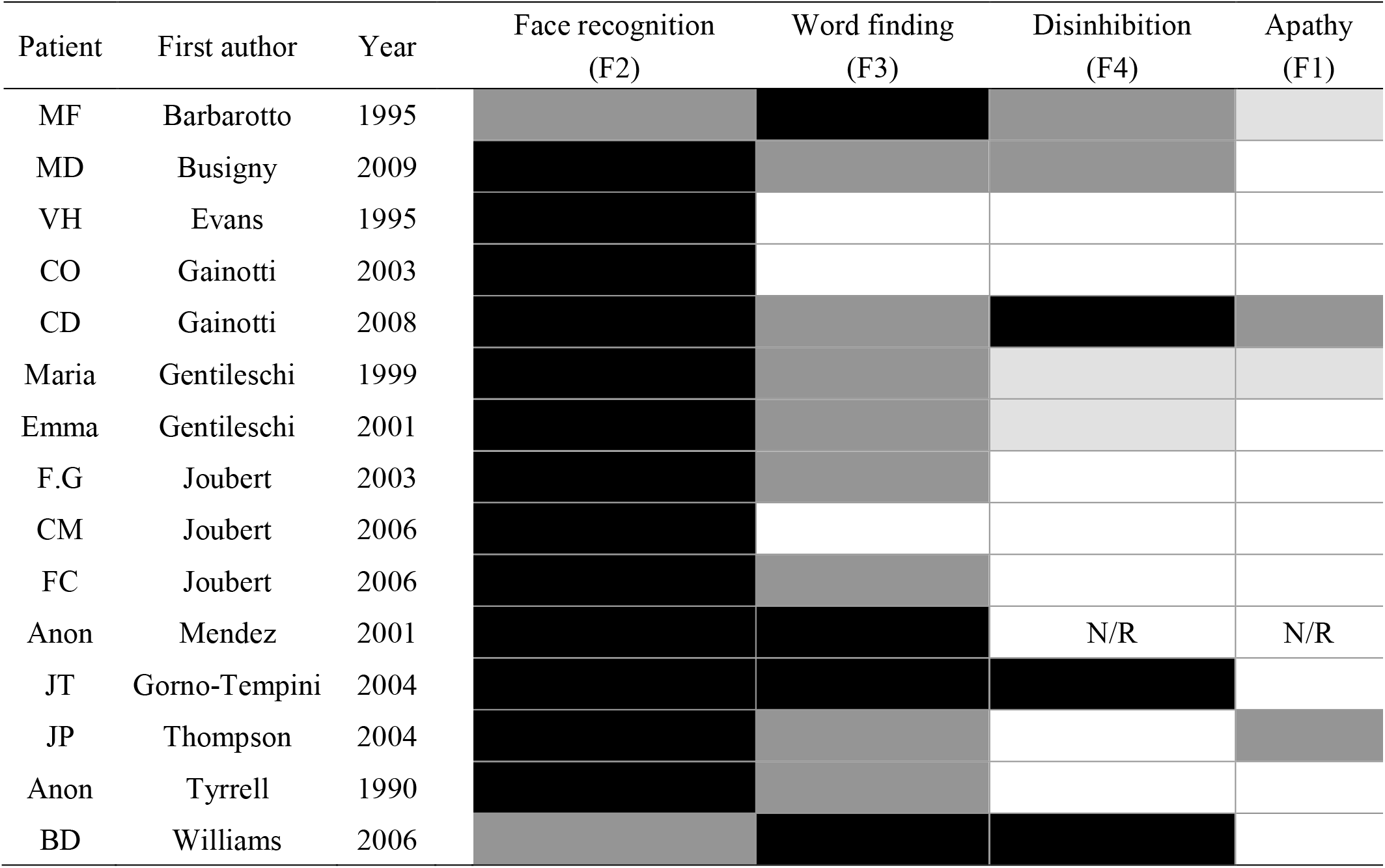
Systematic review of right-sided temporal lobe atrophy cases

## Discussion

The purpose of this investigation was to generate, for the first time, a unified model of ATL functions which captures the known cognitive-behavioural variations in semantic dementia, maps these to the underlying patterns of atrophy, and integrates with the considerable database on healthy semantic function and semantic impairment in other patient groups (for a recent review, see Lambon Ralph et al^23^). This aim was achieved through the analysis of a large SD cases series that varied in terms of severity and the balance of left vs. right ATL atrophy (i.e., representing the typical distribution of cases^40^). Rather than forcing a categorical framework onto the continuous variations across the patients, the new analytical approach was able to capture the graded neuropsychological differences and map these to the patients’ distribution of frontotemporal atrophy. As expected from previous descriptions of SD^2^, the multiple regression analyses confirmed that the degree of generalised semantic impairment (across various verbal and nonverbal semantic tasks) was related to the patients’ total, bilateral ATL atrophy. Verbal production and word-finding abilities were also related to total ATL atrophy as well as to the balance of left>right ATL atrophy – reflecting the enhanced importance of left ATL regions to speech production (see below). Behavioural apathy was found to relate positively to the degree of orbitofrontal atrophy and negatively to total temporal volumes (i.e., those patients with a more frontotemporal balance of atrophy in the context of a SD temporal-variant, selected group). The data-driven PCA replicated and extended these findings by identifying five statistically independent behavioural factors and their unique atrophy correlates. A generalised severity factor was related to increased atrophy around the perimeter of the frontotemporal regions implicated in SD. Again, naming was uniquely correlated with the degree of left ATL atrophy and apathy to medial OFC volumes. In addition, disinhibited behaviour was uniquely correlated with right dorsal STG and OFC atrophy and face recognition to right ATL volumes.

Before briefly reviewing how each of these findings relates to the broader clinical and cognitive neuroscience literature, it is important to consider what these PCA dimensions mean. Although it might be tempting to think of each dimension as a clinical subgroup, this is not correct. By definition PCA and other similar statistical techniques attempt to find continuous, graded dimensions underpinning the variations in the observed data, rather than identify clusters of cases. In fact there is an example of this in the current study; although, like most clinical SD samples^40^, there were no patients with early right-ATL only atrophy, the PCA was able to reveal that (a) there was independent variation in the degree of face recognition deficits and (b) in turn, this was related to the degree of right-ATL atrophy. A more appropriate analogy for the PCA dimensions would be the ingredients in cooking or baking. The variation in patient presentations is like the array of breads, cakes and patisseries found in a bakery, with this diversity reflecting the differing amounts of the key ingredients (flour, butter, eggs, etc.). In effect, the multiple regression and PCA analyses offer a unified model of semantic dementia by identifying and quantifying the ‘ingredients’ that underpin the patients’ clinical variations (i.e., semantic impairment, anomia, prosopagnosia, disinhibited behaviour and apathy). In turn, when these factors are related to the distribution of atrophy, the resultant maps show regions that are uniquely related to each factor (areas that contribute to more than one function are not identified through this method but are by the regression analyses).

The paradigmatic symptom of SD is their selective yet progressive, multimodal semantic impairment^2, 3^. This central symptom was found to relate to the degree of bilateral ATL atrophy. This finding aligns closely with data from healthy participants, other patient groups as well as comparative neurosurgical studies (for reviews, see Lambon Ralph et al and Rice et al^23, 74^). For example, distortion-corrected/avoiding fMRI identifies bilateral ATL activations when healthy participants complete various kinds of semantic task^8, 64^, rTMS to left or right ATL generates a transient, selective semantic impairment^7, 46, 86^, and patients with unilateral ATL resection for temporal lobe epilepsy have a mild semantic impairment albeit much less pronounced than that observed in SD^15, 87^. Indeed, the difference between unilateral and bilateral damage was first demonstrated in non-human primates^43, 44^ and one human case^45^ – with initial unilateral resection generating a transient multimodal semantic impairment leading to a considerable, chronic deficit after bilateral removal. Thus, it would appear that the two ATLs work in concert to generate a robust semantic system which is only significantly compromised in bilateral diseases. This bilateral hypothesis has been recently supported by formal computational models of a bilateral semantic system^20^ as well as combined rTMS-fMRI explorations (which show that after left ATL rTMS in healthy participants, there is both upregulation of activity in the right ATL and increased intrahemispheric functional connectivity)^64, 88^.

The second ‘ingredient’ of semantic dementia is anomia, which is probably the most common, presenting symptom. In keeping with previous explorations^17, 89, 90^, the patients’ anomia was found not only to be related to the degree of general semantic impairment (bilateral ATL volume) but also strongly to the left>right ATL atrophy. Again, this result aligns directly with convergent data from fMRI and rTMS in healthy participants^22, 51^ and from patients with unilateral ATL damage^15, 49, 87, 91^ – all pointing to a greater role of left than right ATL in semantically-initiated speech production/naming. Formal computational models have shown how this form of asymmetric involvement in naming can arise from an inherently bilateral ATL semantic system. Specifically, differential connectivity to the left hemisphere prefrontal speech output systems means that the left ATL component becomes especially important in driving speech from semantic input^17, 20, 92^.

The face-recognition-right ATL dimension may reflect a complementary effect of differential connectivity, this time with respect to input to the ATLs. Semantic knowledge about people was found to pattern with general semantic knowledge and was associated with the degree of bilateral ATL damage. In contrast the degree of right ATL atrophy was linked with visual face recognition per se (cf. the classical definition of visual prosopagnosia)^90, 93^. Again, this finding aligns with the results from patients with unilateral right ATL resection who have also been shown to demonstrate greater deficits of visual recognition of familiar people^87, 94^. Furthermore, the progression of rare, early right ATL semantic dementia patients (see Table 4) fits with this result; in the very earliest stage (which is long before most right>left SD present to clinic), right-only SD patients are reported to have visual prosopagnosia (i.e., poor recognition from faces with good semantic knowledge of the same people when probed from another input modality) which later develops into a generalised semantic impairment and anomia, presumably when bilateral ATL pathology has begun to evolve. It is possible that the differential right>left ATL involvement in face recognition may again reflect differential connectivity^22, 87^. It is well established that there is a strong asymmetry in ventral occipital-temporal regions for different visual objects, with face processing exhibiting a rightward bias (the face-form area^95^) and the opposite for word recognition (the visual word-form area^96^). Accordingly, extending the logic and computational demonstrations for the effect of differential connectivity on function^17, 20, 73^, atrophy of right ATL regions might impact much earlier than left ATL atrophy on face recognition because there is strongest visual face input to this part of the bilaterally-distributed semantic system. It is also possible, that, atrophy extends posteriorly into the FFA and other parts of the right-lateralised extended face processing network^97–100^. In our further analysis, the face deficits of SD were found to be correlated with the level of atrophy in all of the nodes in this network.

The two remaining dimensions relate to the behavioural changes observed in SD and FTD patients more generally^71, 101^. The level of disinhibited behaviour (i.e. disinhibition, irritability and agitation) was found to be related to both right dorsal STG and OFC volumes. This result aligns with findings from three other literatures (for the literature review of right dorsal STG, see Table 5 & Figure 4): (a) previous studies have also associated disinhibited behaviour with the right>left SD patients^58–60, 102^, entire FTD and behavioural variant FTD cohorts^71, 101, 103, 104^; (b) consistent with this proposal, *in vivo* human MRI studies have shown that the right dorsal STG and OFC are activated when normal subjects proceesing social concepts information^62–64^. (c) there is direct evidence to suggest that the right dorsal STG and OFC plays a crucial role in a disinhibition network and connects with other regions relaetd to disinhibtion processing^105–108^. One benefit of using the regression and PCA methods is that it is possible to unpick the covariation of atrophy across frontotemporal regions in SD patients, and thus localise more precisely to the small area of right dorsal STG and OFC compared with the widespread face-related area of the right temporal lobe. Consistent with this hypothesis, more early right SD cases (see Table 4) are reported to behave entirely appropriately on disinhibition, though behavioural deficits can be found in some cases, whose atrophy may start from the superior part of the right dorsal STG.

**Figure 4.**
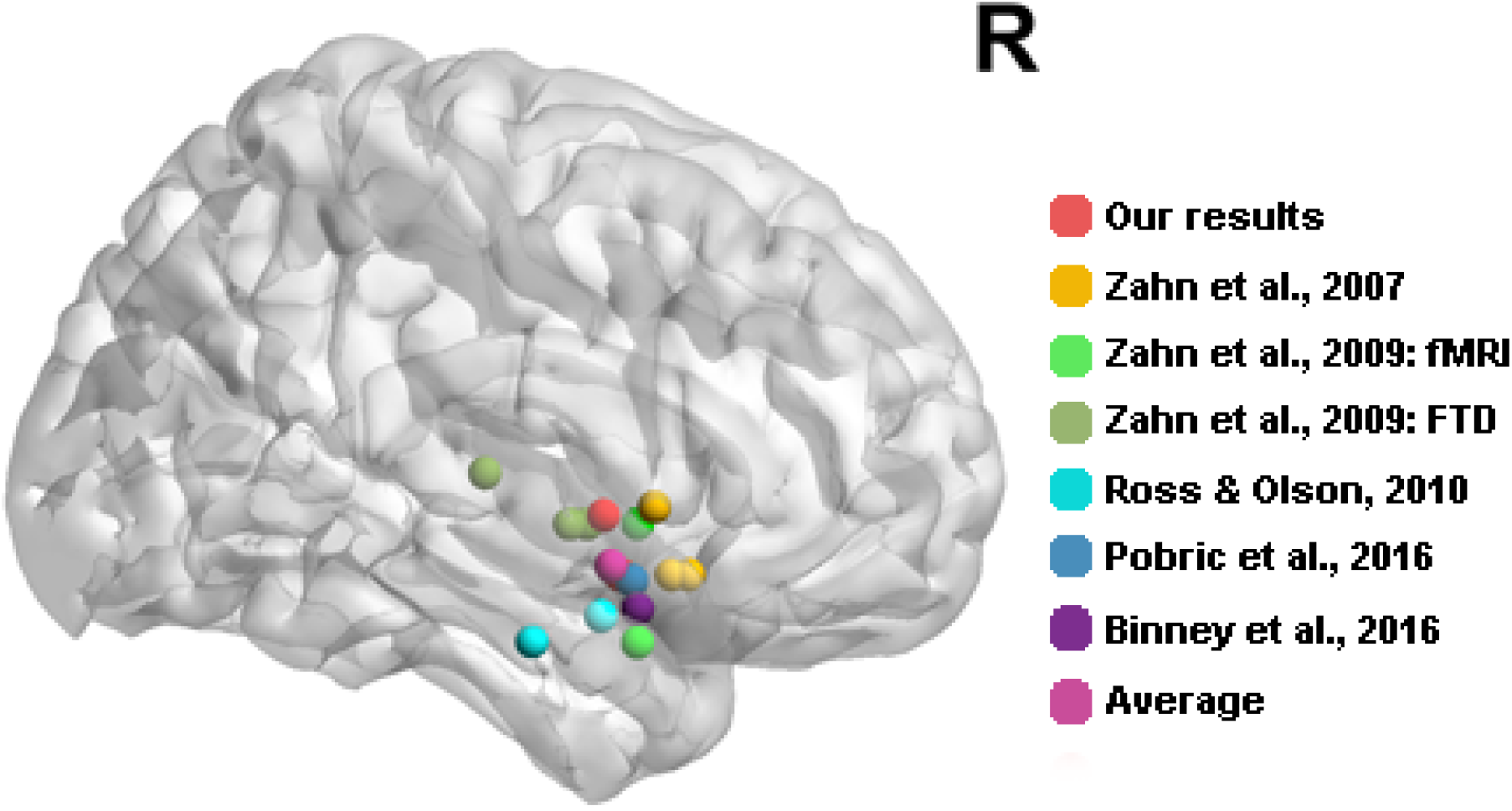
Review of the location of right dorsal STG for social concepts processing from related literature and our study.

**Table 5.** Review of the role of right dorsal ATL in social concepts processing

| First author | Year | Coordinates | Methods | Contrasts |
| --- | --- | --- | --- | --- |
| **Zahn** | 2007 | 57 12 0 | fMRI | Social vs animal function concepts |
| **Zahn** | 2007 | 51 18 -12 | fMRI | Social vs animal function concepts corrected by social concepts vs fixation |
| **Zahn** | 2007 | 51 15 -12 | fMRI | Conjunction of social vs animal function concepts, correlations with descriptiveness of social behaviour and meaning relatedness |
| **Zahn** | 2009 | 54 0 -3 | fMRI | Effect of descriptiveness of social behaviour |
| **Zahn** | 2009 | 57 -18 6 | fMRI | Effect of descriptiveness of social behaviour |
| **Zahn** | 2009 | 60 -3 -3 | fMRI | Effect of descriptiveness of social behaviour masked by the same effect in Zahn et al., 2007 |
| **Zahn** | 2009 | 54 9 -24 | FTD | Hypometabolism of FTD with R sup ATL lesion |
| **Zahn** | 2009 | 51 9 -3 | FTD | Hypometabolism of FTD with R sup ATL lesion masked by Hypometabolism of FTD with social concept selective impairment |
| **Ross** | 2010 | 59 3 -19 | fMRI | Social vs neutral car video |
| **Ross** | 2010 | 66 -10 -24 | fMRI | Social vs animal function concepts |
| **Pobric** | 2016 | 53 8 -13 | TMS | Social vs non-social concepts |
| **Binney** | 2016 | 57 9 -18 | fMRI | Social vs matched-abstract concepts |
| **Average** |  | 56 4 -10 |  |  |
| **Our study** |  | 59 3 -2 | SD | Correlation of disinhibition and atrophy |
| **Our study** |  | 51 6 -12 | SD | Correlation of disinhibition and atrophy |

2 FTD: frontotemporal dementia; TMS: Transcranial magnetic stimulation; SD: semantic dementia;

The final dimension related to negative behaviours (i.e. apathy, depression and anxiety). Apathy is a principal symptom of behavioural variant FTD^103, 109^. As a variant of FTD, SD patients can also show apathy but typically less often than behavioural variant FTD^4, 104^. We found that this factor was associated with the patients’ OFC volume and negatively with atrophy in the ATLs – i.e., those SD patients who had relatively more OFC than ATL damage. This aligns directly with previous studies which have shown a relationship between apathy and OFC in FTD patients^71, 103^. We note that some previous investigations have observed more severe apathy in right>left SD patients^102^. Presumably this might reflect the fact that, as found in the current clinical sample, right>left SD patients tend to have more atrophy of the ATL and OFC overall^60, 68, 84^.

## Methods

### Participants

Forty-seven dementia patients with prominent language problems (25 men, 22 women; age: M = 63 years, s.d. = 7 years; range = 46-74 years; education level: M = 12 years; s.d. = 3 years, range = 3-18 years; years from onset: M = 3 years, s.d. = 2 years, range = 1-10 years; MMSE: M = 22, s.d. = 4; range = 13-29) were recruited from Huashan Hospital, Shanghai. They all met the diagnostic criteria for semantic variant PPA^35^. According to the criteria, patients must present marked naming and single-word comprehension problems. Moreover, at least three of the symptoms should be observed: the deficit of object knowledge, surface dyslexia, spared repetition and spared speech production. The exclusion criteria included a history of head trauma, neurological or psychiatric illness and a severe visuoperceptual impairment. The structural MRI was used to support the diagnosis and exclude other potential comorbid conditions such as nondegenerative brain damage. All the patients presented atrophy in the ATL.

Twenty matched healthy participants (8 men, 12 women; age: M = 61 years, s.d. = 4 years; range = 51-69 years; education level: M = 10 years, s.d. = 3 years; range = 2-16 years) were selected as controls. They performed normally on the MMSE (28 ± 1) with no history of the neurological or psychiatric disorder.

All participants were right-handed native Chinese speakers, had normal or corrected-to-normal hearing and vision, and gave written informed consent. The study was approved by the Institutional Review Board of the State Key Laboratory of Cognitive Neuroscience and Learning, Beijing Normal University.

### Assessments

#### Oral picture naming

Participants were instructed to provide the names of photographs presented on the screen. Object naming consisted of five categories (animals, non-manipulable objects, arbitrary artefacts, fruit and vegetables, and tools), each containing 20 items. The coloured photographs were used and the word frequency of items was matched between categories. Face naming was identical to the object naming, except only one category (20 photographs of famous faces) was presented. All the photographs of famous people were black-and-white. The famous people were Chinese actresses or actors, politicians, and players.

#### Word picture verification

In each trial, a picture and word were presented on the screen and participants were asked to judge whether they were the same object or famous person. In one condition the picture and word matched. In the other, the word was replaced with the name of another exemplar from the same category. The match and mismatch conditions were counterbalanced across two assessment sessions with each item only appearing once per session. Only if participants answered correctly in both conditions (i.e., accepting the correct name and rejecting the semantic foil) was the item scored as correct. The stimuli and categories used in this test were the same as for picture naming, but there were only 10 items per category. The word frequency of items was also matched between five object categories.

#### Picture associative matching

This task had the same format as the Pyramid and Palm Trees test ^110^, in which triplets of pictures were presented on the screen. Participants needed to choose which object/face was more semantically associated with one of the two choices. The stimuli and categories used in this test were the same as for picture naming. This test contained 60 trials, each category containing 10 trials. The word frequency of the targets was matched between categories and the word frequency of answers and foils were matched within each object category.

#### Visual perception for objects and faces

Object perception task included 25 items, each in an array of three line drawings. The target was accompanied by two pictures from a different view. Participants were asked to match the pictures of the same object. The pictures in each trial were from the same category. Similarly, the face perception test required participants to judge whether two faces from different views were from the same person. Thirty-six items were included in this assessment. All the pictures were black-and-white photographs of male faces.

All the above assessments were from our home-made semantic battery but their psychometric properties have been systemically examined (e.g. sensitivity and specificity) among Chinese population. Moreover, they have been widely used in our previous studies about semantic processing on dementia, stroke patients and normal controls^111–113^. Assessments were administered using DMDX ^114^ in separate sessions in a fixed order to avoid the influence of words to pictures. Each session lasted no more than 2 hours. Rest breaks were allowed (see Ding et al^115^ for details). All participants completed this test battery, except one patient who did not finish the word-picture verification. Comparisons between controls and patients, left and right SD patients were assessed using independent two-sample *t*-tests.

#### Behavioural assessments

The neuropsychiatric inventory questionnaire (NPI-Q)^116^ was completed by the caregivers of 24 patients. They rated the presence and severity of twelve behaviour symptoms. Scores range from 0 to 3, representing absent, mild, moderate and severe changes, respectively. This version was the brief form of the NPI and its psychometric properties have been validated^117^. Because some symptoms included in the NPI were rarely endorsed for this patient group, we only chose six symptoms which occurred frequently. Therefore, symptoms of agitation, depression, anxiety, apathy, disinhibition and irritability were entered into our analyses. Chi-square tests were performed to compare the symptom severity between left and right SD groups. The NPI scores were transformed into negative z scores, so that both the cognitive and behavioural scores ran in the same direction (low scores representing poor performance).

### MRI

#### Parameters

The 3D T1 images of all participants were collected through the Siemens 3T scanner. The parameters were listed as follows: repetition time = 2300 ms, echo time = 2.98 ms, flip angle = 9°, matrix size = 240 * 256, field of view = 240 mm * 256 mm, slice number = 192 slices, slice thickness = 1 mm and voxel size = 1 mm * 1 mm * 1 mm.

#### Preprocessing

The images were preprocessed with unified segmentation of SPM 12 (http://www.fil.ion.ucl.ac.uk/spm/). First, bias regularisation was conducted to remove the intensity inhomogeniety caused by the physics of MR scanning (light regularisation; bias FWHM = 60mm). Then the images were segmented into grey matter (the number of Gaussians = 2), white matter (the number of Gaussians = 2), cerebrospinal fluid compartments (CSF; the number of Gaussians = 2), bone (the number of Gaussians = 3), soft tissue (the number of Gaussians = 4) and background (the number of Gaussians = 2) according to the tissue probability map of SPM. Next, they were normalized into the MNI space with both affine and non-linear transformations, resampled to 1.5 * 1.5 * 1.5 mm and modulated to compensate for the image warping during normalization. Finally, they were smoothed with 8mm FWHM. The total intracranial volume was calculated by summing the normalized grey matter, white matter and CSF images together.

### The atrophy of SD patients

#### Voxel-based analysis

Voxel-based comparisons were employed between SD groups and controls (voxel-wise FDR corrected *p* < 0.05). The voxels remaining in SD vs. controls were binarized as a mask for further voxel-based analyses.

#### ROI-based analysis

As a variant of frontotemporal dementia, SD patients showed widespread frontotemporal atrophy^118^. To characterize the atrophy degree of temporal and frontal lobes respectively, four ROIs were derived from the Harvard-Oxford Atlas^119^ (see Figure 1d). The bilateral ATL ROIs consisted of the anterior fusiform gyrus, anterior inferior temporal gyrus and temporal pole. Meanwhile, the bilateral OFC ROIs were composed of the orbital frontal and subcallosal cortex.

Three ROI measures were calculated. The difference between left and right ATL atrophy was generated by comparing the grey matter volumes of left ATL minus right ATL. The sums of ATL regions across hemispheres were used as the total ATL atrophy measure and OFC regions for total OFC atrophy. Independent two-sample t-tests were implemented separately between SD groups vs. normal controls, and left vs. right sided SD patients. In addition, the z-scores of patient’s atrophy status across voxels was generated by regularizing their grey matter volumes with respect to the control cohorts. The corresponding z-scores for the three ROI measures were extracted for subsequent analyses. For the initial analyses that split patients according to the balance of temporal lobe atrophy, patients were divided into left>right and right>left according to the [left – right ATL] ROI measure.

### The relationship between atrophy and task performance

To understand how the laterality of temporal lobe atrophy, the total bilateral temporal or frontal lobe atrophy influenced patients’ task performance, we built a series of general linear models, which used the three ROI z-scores as independent variables to predict all eight cognitive task measures and six NPI items.

### The PCA of task performance

To explore the underlying dimensions of variation in the patients’ cognitive and behavioural measures, PCA analysis was implemented on the three face semantic, three object semantic, two visual perceptual tasks and six NPI items. Some subjects’ data were missed, so we imputed their data to increase our statistical power for our PCA-related analysis. To achieve this, we first used the function ‘pca_compsel’ of PCA toolbox (http://michem.disat.unimib.it/chm/) to determine the number of optimal factors of our data. Then the missing data were imputed by the function ‘ppca’ of matlab (https://www.mathworks.com/products/matlab.html). Finally, the imputed data were entered into the PCA by JMP (https://www.jmp.com/en_us/home.html). All the scores were converted to z-scores before PCA. The factors with eigenvalue > 1 were extracted and varimax rotation applied to enhance cognitive interpretability of the principal components.

### The relationship between atrophy and PCA factors

#### ROI-based Regression

To identify the relationship of temporal and frontal lobe status to the PCA factors, the corresponding ROI-based variables were again entered into the regression models using the same method as for the regions models of specific task data (see above). The only exception here was that the dependent task variables were replaced by PCA factor scores.

#### Voxel-based correlational methodology (VBCM)

To explore whether other regions beyond the ROIs also associated with the patients’ PCA scores, we completed voxel-based correlation analyses between the voxel-based z scores and each PCA factor within the atrophy mask. Due to the wide range of statistical powers, different voxel-wise thresholds were adopted for the factors (voxel *p* < 0.001, *p* < 0.01 or *p* < 0.05; cluster size > 50 voxels).

### Face ROI analysis

Face processing is associated with widespread right temporal areas ^97^, thus we explored which regions were involved in SD’s face deficits. Five classic face-related ROIs were chosen from previous literature (the Talairach coordinates of occipital face area: 25, −88, −10; fusiform face area: 40, −44, −16; posterior superior temporal sulcus: 48, −48, 8; ventral ATL: 25, 0, −28; anterior superior temporal sulcus: 48, −12, −7)^120^. We transformed these Talarich coordinates into the MNI space and generated 6-mm spheres for all ROIs. Then, we extracted the z scores of these ROIs and correlated them with the face-related PCA scores. To validate the specific role of these right-sided ROIs in face processing (over the left hemisphere homologues), supplementary analyses were conducted to (1) correlate the right ROI z scores with object processing tasks; (2) correlate the left corresponding ROI status with face processing scores; and (3) correlate the right ROI volumes in the control group against face processing performance.

## Acknowledgements

This work was supported by the Chinese Scholarship Council to JD & HL; Medical Research Council programme grant (MR/R023883/1) and ERC grant (GAP: 670428 - BRAIN2MIND_NEUROCOMP) to MALR; National Natural Science Foundation of China (31872785 & 81171019) and Beijing Natural Science Foundation (7182088) to ZH; National Key R&D Program of China (2016YFC1306305) to QG.

## Author contributions

The study idea was proposed and supervised by MALR, QG and ZH. The behavioural and imaging data were collected by KC, QY, LH and YL, respectively. The behaviour comparison and PCA were carried out by HL and YC. The brain-behaviour analysis was carried out by JD and KC. The paper was written by MALR and JD. The figures and tables were made by MALR, JD and HL.

