## Supplementary materials for "A unified neurocognitive model of the anterior temporal lobe contributions to semantics, language, social behaviour & face recognition"

Supplementary table 1. The correlation values of right face related ROIs in SD patients.

| | OFA | FFA | pSTS | vATL | aSTS |
|---|---|---|---|---|---|
| **Face** | | | | | |
| Word picture verification | 0.27 | **0.43**** | **0.30*** | **0.46**** | **0.55***** |
| Picture matching | 0.27 | **0.36*** | 0.27 | **0.42**** | **0.39**** |
| Picture naming | 0.12 | 0.09 | 0.00 | -0.01 | 0.05 |
| Perception | **0.37*** | 0.24 | **0.37*** | **0.39**** | **0.42**** |
| **Object** | | | | | |
| Word picture verification | 0.07 | 0.06 | -0.02 | 0.04 | 0.19 |
| Picture matching | 0.01 | 0.22 | 0.05 | **0.38**** | **0.35*** |
| Picture naming | -0.07 | -0.02 | -0.13 | -0.16 | -0.07 |
| Perception | 0.16 | -0.13 | 0.09 | -0.24 | -0.19 |

*: $p < 0.05$; **: $p < 0.01$; ***: $p < 0.001$.

OFA: occipital face area; FFA: fusiform face area; STS: superior temporal sulcus; ATL: anterior temporal lobe.

Supplementary table 2. The correlation values of left face related ROIs in SD patients.

| | OFA | FFA | pSTS | vATL | aSTS |
|---|---|---|---|---|---|
| **Face** | | | | | |
| Word picture verification | 0.14 | 0.18 | 0.25 | **0.35*** | 0.22 |
| Picture matching | 0.09 | **0.33*** | **0.35*** | **0.43**** | **0.34*** |
| Picture naming | 0.04 | 0.25 | 0.13 | **0.29*** | **0.33*** |
| Perception | 0.23 | 0.09 | 0.19 | **0.30*** | 0.03 |
| **Object** | | | | | |
| Word picture verification | 0.03 | **0.30*** | 0.19 | **0.44**** | **0.43**** |
| Picture matching | -0.01 | 0.19 | 0.10 | **0.43**** | 0.17 |
| Picture naming | 0.01 | **0.47***** | 0.21 | **0.52***** | **0.52***** |
| Perception | 0.08 | 0.13 | 0.18 | 0.13 | 0.11 |

*: $p < 0.05$; **: $p < 0.01$; ***: $p < 0.001$.

OFA: occipital face area; FFA: fusiform face area; STS: superior temporal sulcus; ATL: anterior temporal lobe.

Supplementary table 3. The correlation values of left and right face related ROIs in normal controls.

| | OFA | FFA | pSTS | vATL | aSTS |
|---|---|---|---|---|---|
| **Face & right ROIs** | | | | | |
| Word picture verification | -0.19 | -0.30 | 0.27 | -0.12 | -0.23 |
| Picture matching | 0.13 | -0.42 | -0.18 | **-0.46*** | -0.32 |
| Picture naming | 0.04 | -0.29 | 0.33 | -0.04 | -0.18 |
| Perception | 0.14 | -0.13 | 0.20 | 0.08 | 0.06 |
| **Face & left ROIs** | | | | | |
| Word picture verification | 0.06 | 0.08 | -0.30 | -0.33 | -0.07 |
| Picture matching | -0.03 | -0.39 | 0.27 | **-0.54*** | -0.13 |
| Picture naming | 0.13 | -0.02 | -0.07 | -0.20 | -0.13 |
| Perception | 0.28 | -0.25 | 0.03 | -0.02 | 0.03 |

OFA: occipital face area; FFA: fusiform face area; STS: superior temporal sulcus; ATL: anterior temporal lobe.

*: $p < 0.05$; **: $p < 0.01$; ***: $p < 0.001$.
